# Time-resolved proximity labeling of protein networks associated with ligand-activated EGFR

**DOI:** 10.1101/2022.01.07.475389

**Authors:** Mireia Perez Verdaguer, Tian Zhang, Joao A. Paulo, Callen Wallace, Simon C. Watkins, Steven P. Gygi, Alexander Sorkin

## Abstract

Ligand binding to the EGF receptor (EGFR) triggers multiple signal transduction processes and promotes endocytosis of the receptor. The mechanisms of EGFR endocytosis and its crosstalk with signaling are poorly understood. Here, we combined peroxidase-catalyzed proximity labeling, isobaric peptide tagging and quantitative mass-spectrometry to define the dynamics of the proximity proteome of ligand-activated EGFR. Using this approach, we identified a network of signaling proteins, which remain associated with the receptor during its internalization and trafficking through the endosomal system. We showed that Trk-fused gene (TFG), a protein known to function at the endoplasmic reticulum exit sites, was enriched in the proximity proteome of EGFR in early/sorting endosomes and localized in these endosomes, and demonstrated that TFG regulates endosomal sorting of EGFR. This study provides a comprehensive resource of time-dependent nanoscale environment of EGFR, thus opening avenues to discovering new regulatory mechanisms of signaling and intracellular trafficking of receptor tyrosine kinases.

## INTRODUCTION

Epidermal growth factor (EGF) receptor (EGFR) belongs to a large family of receptor tyrosine kinases which control a broad range of cell activities, such as cell proliferation, survival, differentiation, motility and metabolism. EGFR plays important roles in mammalian development and tissue homeostasis in adult organisms (Sibilia et al., 2007). Mutations and overexpression of EGFR often leading to its aberrant activity are associated with tumorigenesis and metastatic processes (Grandis and Sok, 2004). Ligand binding to EGFR triggers receptor dimerization, activation of the tyrosine kinase in its cytoplasmic domain and phosphorylation of tyrosine residues in the receptor carboxy-terminal tail. Phosphotyrosine motifs serve as docking sites for downstream signaling proteins containing Src homology 2 (SH2) and phosphotyrosine binding domains (Lemmon and Schlessinger, 2010). Ligand binding also results in rapid endocytosis through clathrin-dependent and -independent pathways (Sorkin and Goh, 2009). Internalized EGFR is capable of recycling back to the cell surface but is also targeted for lysosomal degradation leading to receptor down-regulation (Sorkin and Goh, 2009). Endocytic trafficking of EGFR is proposed to play an important role in the spatiotemporal regulation of EGFR signaling, and dysregulation of this trafficking is associated with cell transformation (Mellman and Yarden, 2013; Sigismund et al., 2021; von Zastrow and Sorkin, 2021). However, mechanisms of EGFR endocytic trafficking and its crosstalks with EGFR signaling are not well understood. In particular, whether EGFR triggers signaling after endocytosis from endosomes and whether such signaling is necessary for proper downstream outcomes is unclear.

Defining the dynamics of EGFR interactome in time and space is crucial for elucidation of the mechanisms of ligand-induced endocytosis of EGFR and how endocytosis regulates signaling. Numerous mass-spectrometry proteomic studies analyzed proteins co-immunoprecipitated with EGFR and proteins phosphorylated in cells upon EGFR activation [examples are (Foerster et al., 2013; Francavilla et al., 2016; Tong et al., 2014)]. These studies generated one of the most extensive interactome and phosphosite database for any signaling receptor. Moreover, compartment-specific interactome of EGFR was analyzed by identifying proteins co-precipitated with EGFR in various subcellular fractions in HeLa cells (Itzhak et al., 2016). However, all above mentioned studies analyzed samples either after detergent solubilization or cell homogenization, procedures that may disrupt transient protein-protein interactions. By contrast, techniques based on proximity labeling by biotin allow detection of proteins located within ~20-nm radius from the bait in the intact cell. In particular, highly efficient proximity labeling with engineered ascorbate peroxidase APEX2 (Lam et al., 2015) combined with quantitative mass-spectrometry allows fine temporal resolution, and has been used for a space- and time-resolved analysis of the interaction networks of agonist-activated G-protein coupled receptors (Lobingier et al., 2017; Paek et al., 2017).

In the present study, we generated cells stably expressing EGFR-APEX2 fusion protein and analyzed proximity proteome of EGFR-APEX2 during first hour of its activation by EGF. The time-dependent abundance of various plasma membrane, endosome, and lysosome resident proteins in the proximity proteome was consistent with the movement of EGFR-APEX2 upon EGF stimulation from the cell surface to early and sorting endosomes, and then to late endosomes and lysosomes. Searching the EGFR-APEX2 proximity proteome for proteins with time-dependent abundance profiles which are similar to those of resident sorting endosome proteins, we identified a network of proteins implicated in signal transduction processes, suggesting their localization in EGFR-containing endosomes. Moreover, this analysis identified a new putative component of the endosome, TFG (Trk-fusion gene). TFG is a cytosolic protein that is located at the endoplasmic reticulum (ER) exit sites (ERES) and involved in COPII-mediated cargo transport from the ER to the ERGIC compartment (Witte et al., 2011). To explore the finding of TFG in proximity to internalized EGFR, we demonstrated the localization of a fraction of TFG in early and sorting endosomes and showed that it regulates ligand-induced degradation of EGFR.

## RESULTS AND DISCUSSION

### Time-resolved proximity labeling defines the dynamics of EGFR interaction networks during ligand-induced internalization and post-endocytic traffic of the receptor

To analyze time-dependent changes in the proximity proteome of ligand-activated EGFR, we generated human colon carcinoma HCT116 and human embryonic kidney HEK293T cell lines that stably express EGFR fused at the C-terminus with an engineered ascorbate peroxidase with higher catalytic activity (APEX2) (Lam et al., 2015) (Figure 1A). Western blotting analysis showed that EGFR-APEX2 is expressed in HCT116 cells at a high level that was comparable with endogenous EGFR, whereas the expression level of EGFR-APEX2 in HEK293T cells is significantly lower (Figure 1B). Figure 1A illustrates the pipeline of a proximity labeling experiment in which serum-starved HCT116/EGFR-APEX2 or HEK293T/EGFR-APEX2 cells are preincubated with biotinyl-tyramide for 1 h and stimulated with 100 ng/ml EGF. Biotinylation of cytoplasm-exposed protein regions located in the proximity to APEX2 (20 nm radius) is catalyzed by freshly diluted H_2_O_2_ (final concentration 1 mM for 1 min). After cell lysis and Streptavidin-Biotin pulldown, precipitated biotinylated proteins are digested by trypsin, and labelled using Tandem Mass Tag (TMT) 9-11plex reagents.

**Figure 1.**
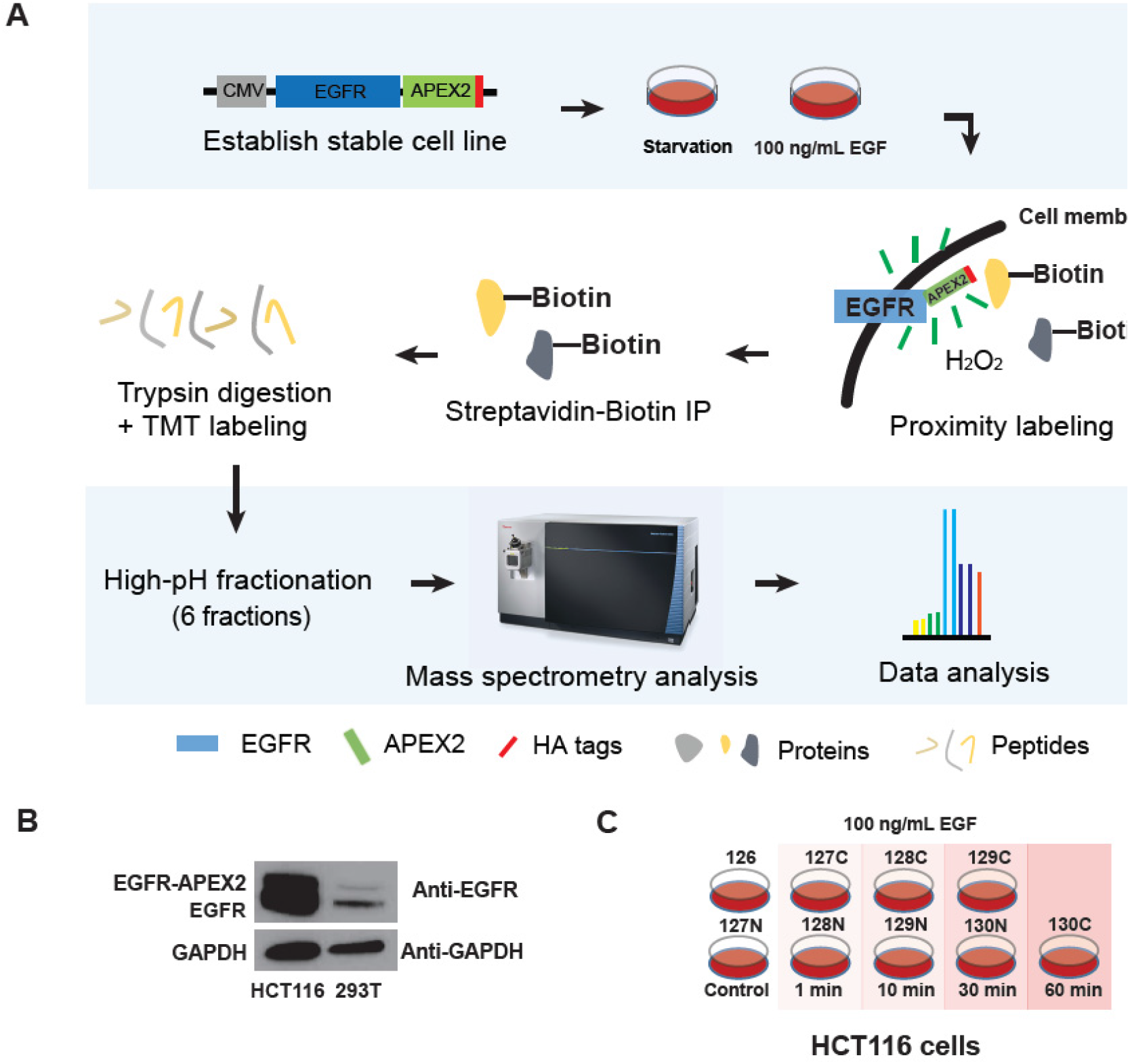
Schematics of proximity labeling experiment of EGFR-APEX2 to study endocytosis. **(A)** HCT116 and HEK293T cells are engineered to stably express EGFR fused with APEX2 tag. Serum-starved cells are stimulated with EGF to activate EGFR-APEX2 and cause its endocytosis. During 1 min of H_2_O_2_ treatment, APEX2 catalyzes the transfer of biotin phenol (BP) to proteins surrounding EGFR-APEX2 within a radius of 20 nm. Biotinylated proteins were pulled down by streptavidin immunoprecipitation, digested, TMT-labelled, combined in a TMT experiment, fractionated into 6 fractions, and analyzed by mass spectrometry. Data analysis reveals the dynamics of relative abundances of proteins with proximity to EGFR following EGF treatment. (**B**) Western blots showing expression of EGFR-APEX2 in HCT116 cells and HEK293T cells. (**C**) Experimental design of proximity labeling TMTplex9 experiment in HCT116 cells presented in Figure 2. Cells after 24 hrs of serum starvation were used as control. Cells were treated using 100 ng/ml EGF for 1-minute, 10-minute, 30-miunte and 60-miunte and collected for time-course analysis.

To track time-dependent proximity proteome of EGFR-APEX2, serum-starved HCT116 cells were left untreated or incubated with EGF for 1, 10, 30 and 60 min. After trypsin digestion, labeling with TMT 9plex was performed (Figure 1C), samples were fractionated into six fractions and analyzed by mass spectrometry (Figure 1A). Heatmap of 1797 proteins quantified in this experiment showed that many proteins had distinct time-dependent proximity to EGFR-APEX2 after EGF treatment (Figure 2A). All quantified proteins were sorted into three clusters by a K-means clustering method. Proximity to EGFR-APEX2 of Cluster 1 proteins did not significantly change after cell stimulation with EGF. Proteins in Cluster 2 displayed decreased proximity to EGFR-APEX2, while proteins in Cluster 3 had increased proximity to EGFR-APEX2 in EGF-stimulated cells (Figure 2A). Gene ontology analysis showed that Cluster 2 was enriched in plasma membrane proteins, while Cluster 3 was enriched in proteins of endosome-related gene ontology categories (Figure 2B). Such dynamics of the proximity proteome is consistent with redistribution of ligand-bound EGFR-APEX2 from the plasma membrane to endosomes.

**Figure 2.**
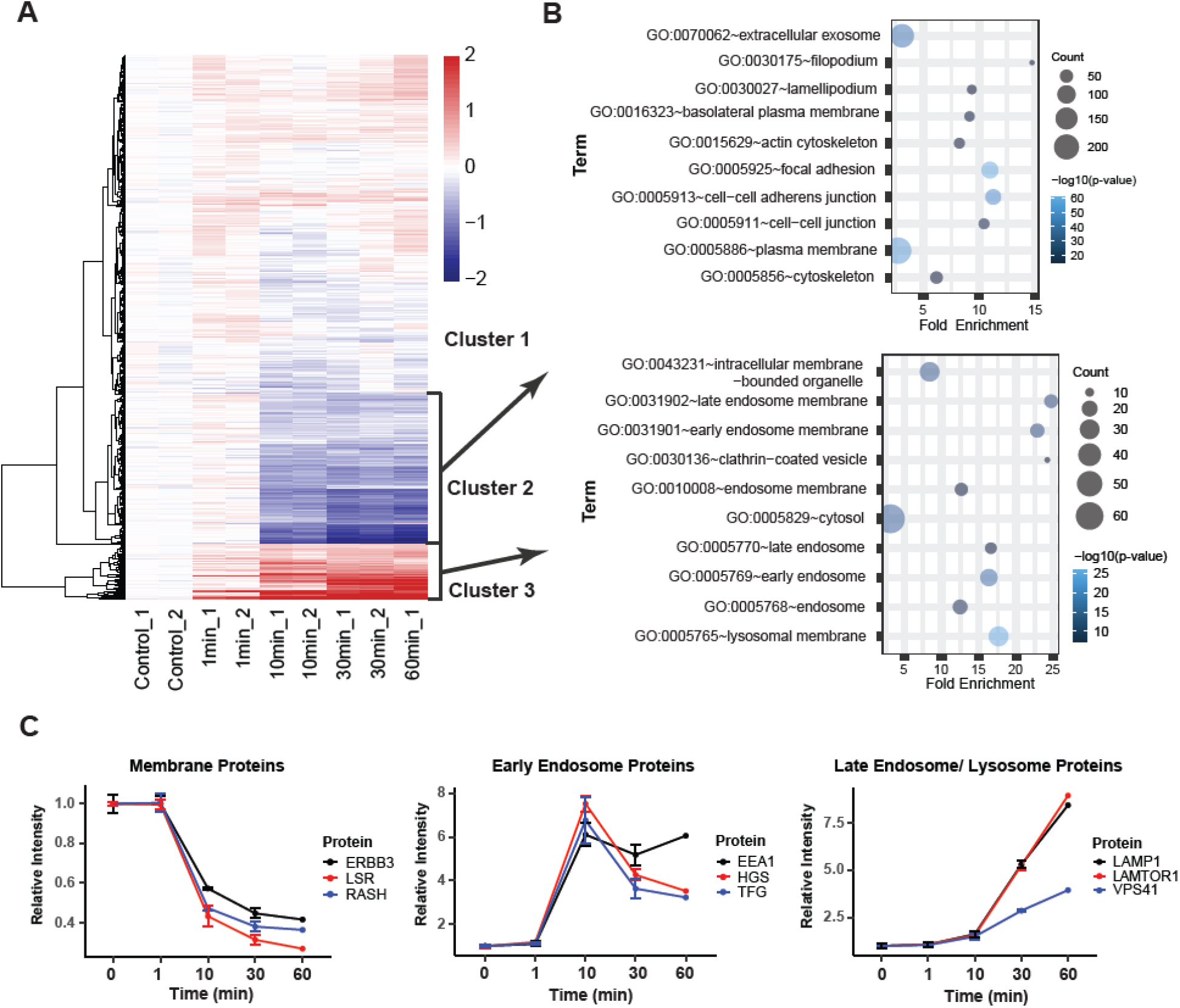
Time-course of relative abundances of EGFR-APEX2 proximal proteins following EGF treatment. (**A**) Heatmap of proteins quantified in APEX2 experiment in HCT116 cells designed as shown in Figure 1C. 1797 proteins with a summed signal-to-noise greater than 100 and variation lower than 20% in each time point are shown. Proteins were sorted into 3 clusters using K-means clustering. (**B**) Gene ontology analysis was conducted with proteins from Cluster 2 and Cluster 3 in the heatmap, respectively. Upper panel shows that Cluster 2 with proteins of lower relative abundance after EGF treatment (Cluster 2) was enriched in plasma membrane proteins. Cluster 3 of proteins with higher relative abundance after EGF treatment was enriched in endosomal proteins. (**C**) Examples of time-dependencies of relative abundances in EGFR-APEX2 proximity of plasma membrane (Cluster 2), early/sorting endosome (Cluster 3) and late endosome/lysosome proteins (Cluster 3) following EGF stimulation.

Examples of the time-dependent abundance of Cluster 2 proteins are presented in Figure 2C. The relative intensity of ErbB3, a member of receptor tyrosine-kinase family and a heterodimerization partner of EGFR, small GTPase HRAS, a downstream signaling effector of EGFR, and a resident plasma membrane protein LSR (lipolysis-stimulated lipoprotein receptor) decreased ~2-times after EGF treatment for 10 min (Figure 2C). Such changes agree with the internalization impairment of ErbB3 (Baulida et al., 1996) and the predominant localization of endogenous HRAS in the plasma membrane (Pinilla-Macua et al., 2016). By contrast, cluster 3 proteins, such as EEA1 (an early endosome-associated antigen 1) and HGS, a hepatocyte growth factor-regulated tyrosine kinase substrate (further mentioned as HRS, a common protein name of *HGS*) (Komada and Kitamura, 1995), which are associated with early and sorting endosomes (Wenzel et al., 2018), had increased proximity to ligand-activated EGFR-APEX2 with the maximum abundance at a 10-min time point (Figure 2C). Time-dependent enrichment profiles in EGFR-APEX2 proximity proteome of several resident early/sorting endosome proteins, such as STAM, STAM2, PTPN23, VPS37A, USP8 and others, were virtually identical to the profile of HRS (Supplemental Table1. The data are available at https://wren.hms.harvard.edu/egfr/). On the other hand, proteins known to reside in late endosomes and lysosomes, such as LAMP1, LAMTOR1 and VPS41, were found to be enriched in the proximity to EGFR-APEX2 after 30-60 min of EGF stimulation (Figure 2C). Together, the data in Figure 2 demonstrate that the dynamics of the time-resolved proximity proteome of EGFR-APEX2 recapitulate ligand-induced endocytosis and post-endocytic trafficking of the receptor through the endolysosomal system and provide comprehensive information about changes in the receptor-proximal protein networks.

Comparative analysis of the EGFR-APEX2 proximity proteome in HCT116 and HEK293T cells incubated or not with 100 ng/ml EGF for 5 min revealed two major clusters of EGFR proximal proteins enriched in EGF-stimulated cells: (i) proteins that are part of the general endosomal sorting machinery; and (ii) proteins that are not residents of endosomes but are known to be involved in signal transduction processes (Figure 3). Examples of proteins enriched in the second, “signaling” cluster of the proximity proteome of EGF-activated EGFR-APEX2 are phosphotyrosine-binding adaptors, such as Grb2 and SHC1, and SH2-domain-containing enzymes, such as RASA1 (GTPase activating protein of RAS) and PLCG1 (phospholipase γ1), all previously shown to remain associated with EGFR in endosomes [reviewed in (Sorkin and Von Zastrow, 2002)] (Figure 3B-C). Importantly, proximity labeling suggested the presence in early/late endosomes of several other signaling proteins that are not known to be directly associated with EGFR or enriched in endosomes. The list of such proteins includes adaptors like GAB1, NCK2, and CRK, GAREM (GRB2-associated and regulator of MAPK protein 1), non-receptor tyrosine kinases, such as TNK2/ACK1 and CSK (kinase upstream of Src family kinases); and cytoskeleton-regulating proteins like WASL (Neural Wiskott-Aldrich syndrome protein; N-WASP) that promotes actin polymerization (Figure 3B and C). Analysis of time-dependent abundance profiles for proteins co-enriched with HRS and other sorting endosome proteins in experiment described in Figure 2 further expands the network of signaling proteins that are proximal to EGFR in endosomes with PIK3R2 and PIK3RCA/B (regulatory and catalytic subunits of phosphotidyl-inositol-3-kinase, PI3K), adaptor/scaffold proteins such as SPRY4, SH3KBP1 and a PDZ domain protein INADL (the data are available at https://wren.hms.harvard.edu/egfr/). While the precise roles of many of these signaling proteins in EGFR-containing endosomes need to be further investigated, the data in Figures 2 and 3 suggest that multiple signaling pathways can be triggered from endosomes containing EGF-activated EGFR. Alternatively, co-internalization of signaling effectors with EGFR may serve to terminate their signaling from the plasma membrane as part of negative feedback regulatory mechanisms.

**Figure 3.**
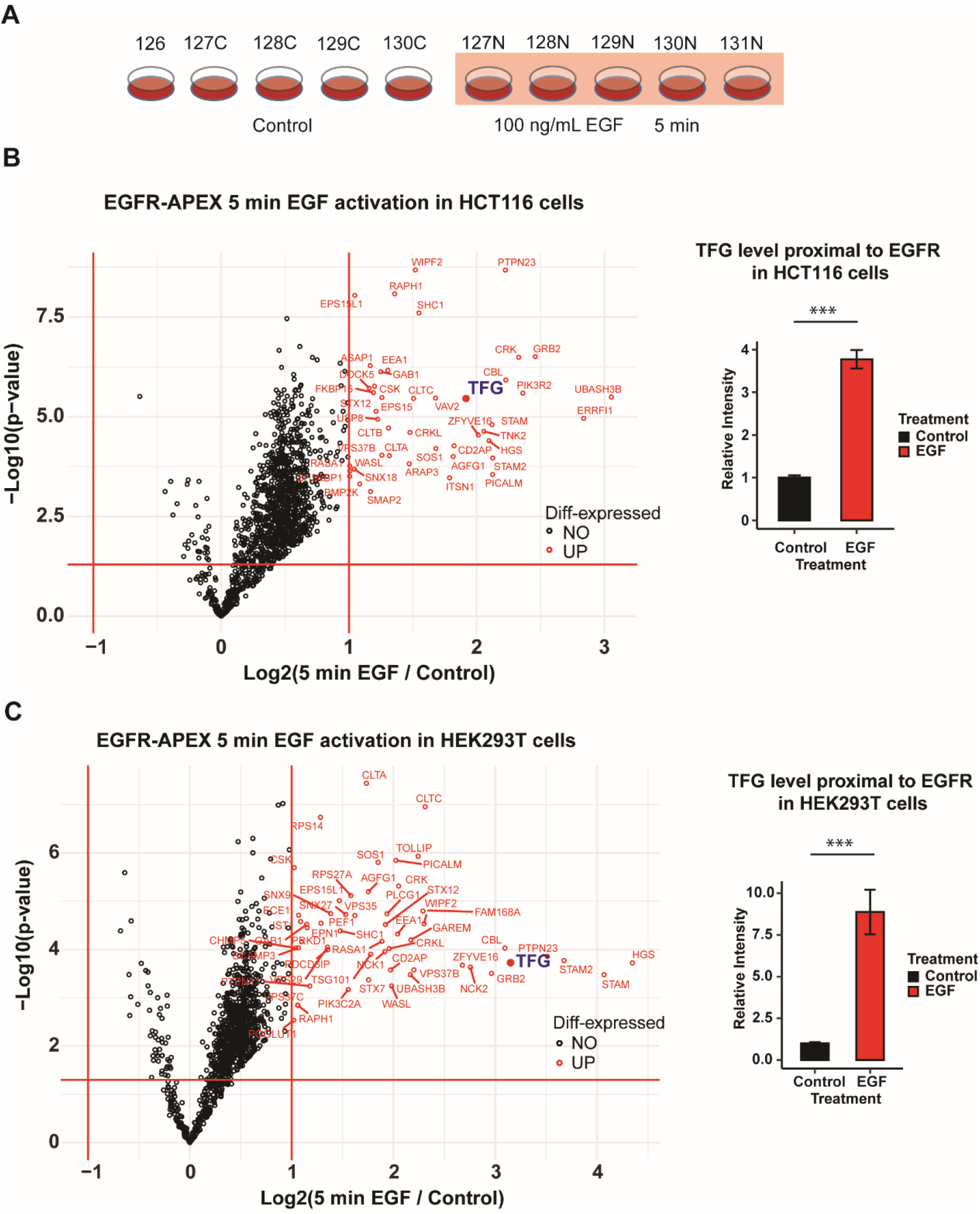
Enrichment of endosomal and signaling proteins in EGFR proximal proteome upon 10-min EGF stimulation. (**A**) Schematics of the proximity labeling TMTplex10experiment of EGFR in HCT116 and HEK293T cells. Serum-starved cells were either untreated (control) or treated with 100 ng/ml EGF for 5 min and used to analyze the proximal proteins. (**B-C**) Volcano plots of quantified proteins in EGFR-APEX2 experiments in HCT116 (**B**) and HEK293T cells (**C**). Proteins with greater than 2-fold change in relative abundance after EGF treatment are highlighted in red. Relative abundance of TGF in EGFR-APEX2 proximity increased significantly in the two cell lines. p-value < 0.001 (graphs on the right).

### TFG is enriched in the proximity proteome of endosomal EGFR-APEX2

Most proteins that were enriched in the proximity proteome of EGFR in early/sorting endosomes (Figures 2 and 3) have an established or a putative function either in endosomal traffic or intracellular signaling. Interestingly, Trk-fused gene (TFG) protein was found in the EGFR-APEX2 proximity proteome with maximum abundance at 10-min of EGF stimulation (Figure 2C). The time-dependent enrichment profile of TFG was virtually identical to that of HRS (Figure 2C) and other proteins of sorting endosomes, such as STAM, STAM2 and PTPN23 in HCT116 cells (Supplemental Table 1). Robust enrichment of TFG was observed in both HCT116 and HEK293T cells stimulated with EGF for 5 min (Figure 3). Finally, time-course of relative protein abundances during first 10 min of EGF stimulation in TMTplex11 experiments (Figure 4A) demonstrated that TFG accumulates in EGFR-proximity proteome with Cluster 3 in both cell lines (Supplemental Figure S1) and that the maximum of TFG enrichment in EGFR proximity proteome was reached earlier in HEK293T cells, presumably due to a substantially lower level of EGFR-APEX2 in these cells than in HSC116 cells (Figure 4A-C). Altogether, APEX2 experiments presented in Figures 2–4 suggest that TFG could be a novel component of sorting endosomes containing EGFR. Previous studies showed that TFG is a polymeric scaffold located at the interface between ERES and the ERGIC compartment and proposed that TFG is involved in uncoating of COPII vesicles and their delivery to the ERGIC-target membrane (Johnson et al., 2015; Witte et al., 2011). However, relative abundances of ERES proteins Sec16, Sec13 and Sec31, all proposed to interact with TFG (Hanna et al., 2017; Huttlin et al., 2021; Witte et al., 2011) did not significantly change upon EGF stimulation (Figures 2C and 4A-B), indicating that these proteins are not enriched in the proximity of endosomal EGFR-APEX2. Therefore, finding of TFG proximity to the cytosolic domain of endosomal EGFR is indicative of ERES-independent localization and function of TFG, which motivated us to focus on microscopic examination of TFG localization in the cell and its endosomal function.

**Figure 4.**
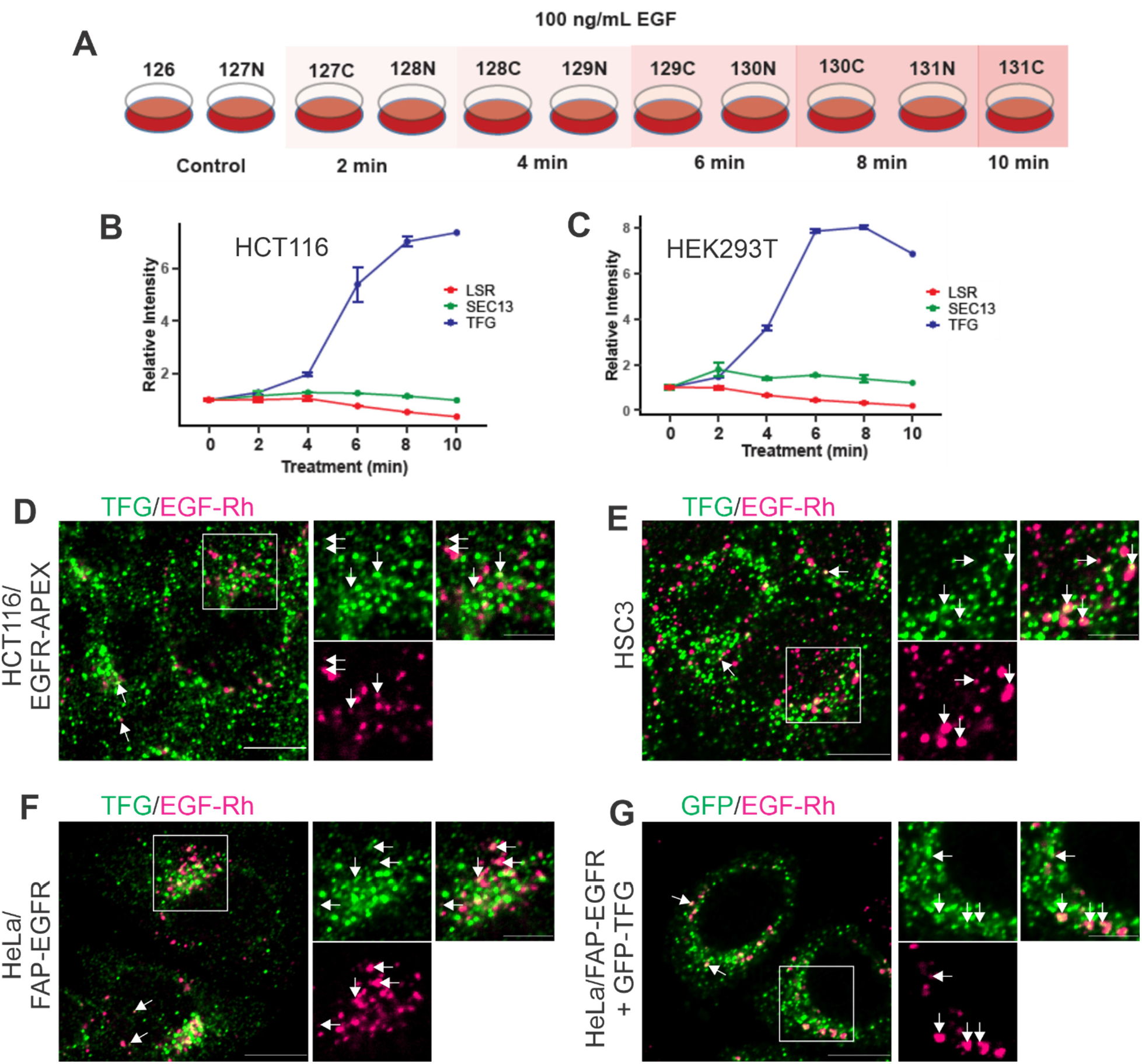
Time-dependent enrichment of TFG in the proximity of EGF-activated EGFR-APEX2 and colocalization of TFG with internalized EGF-Rh. (**A**) Schematics of the proximity labeling TMT plex11 experiment in HCT116 and HEK293T cells. Serum-starved cells were either untreated (control) or treated with 100 ng/ml EGF for 2-10 min to analyze the proximal proteins during receptor internalization. Replicates were included except for 10 min treatment. (**B-C**) Time course of the relative intensity of TGF compared to LSR (plasma membrane) and Sec13 (ERES) proximal to EGFR-APEX2 in HCT116 (***B***) and HEK293T (***C***) cells stimulated with EGF in the experiment described in (***A***). (**D**-**F**) HCT116/EGFR-APEX (**D**), HSC3 (**E**) and HeLa/FAP-EGFR (**F**) cells were stimulated with 10 ng/ml EGF-Rh for 15 min, fixed and stained with TFG antibodies. 3D confocal images were acquired through 561 nm (*magenta*, EGF-Rh) and 488 nm channels (*green*, TFG). (**G**) HeLa/FAP-EGFR cells transfected with GFP-TFG were stimulated with 10 ng/ml EGF-Rh for 15 min. 3D confocal images were acquired through 561 nm (*magenta*, EGF-Rh) and 488 nm channels (*green*, GFP-TFG). Insets in ***D-G*** represent enlargements of the regions indicated by rectangles. Single sections in the middle of the cell of representative images are shown. Arrows indicate examples of colocalization. Scale bars, 10 μm in full images and 5 μm in insets.

### TFG partially co-localizes with internalized ligand-bound EGFR

Immunofluorescence microscopy of HCT116/EGFR-APEX2 cells, that were stimulated with EGF-rhodamine conjugate (EGF-Rh) for 15 min and stained with the TFG antibody, revealed a punctate distribution pattern of endogenous TFG (Figure 4D). A fraction of TFG puncta colocalized with internalized EGF-Rh (Figure 4D), thus supporting the observation of TFG proximity to endosomal EGFR discovered in EGFR-APEX2 labeling experiments. To further characterize endosomal localization and function of TFG in cells expressing endogenous EGFR, we used human squamous carcinoma HSC3 cells, which are growth-dependent on EGFR (Kudo et al., 2003), and HeLa cells expressing endogenous EGFR tagged with fluorogen activating protein (FAP) by gene-editing, which we thoroughly characterized as EGFR endocytosis model system (Larsen et al., 2019). As shown in Figures 4E and F, TFG was partially co-localized with internalized EGF-Rh in both these cell lines as well. Furthermore, GFP-tagged TFG (GFP-TFG) transiently expressed in HeLa/FAP-EGFR was readily detected in EGF-Rh containing endosomes (Figure 4G).

To compare the extent of TFG localization at ERES and in endosomes, HSC3 cells were costained with antibodies to TFG and Sec31, a component of COPII coat that is involved in the ER protein export at ERES (Peotter et al., 2019). Figures 5A and G shows that, in agreement with previous studies (Hanna et al., 2017; Johnson et al., 2015), most bright TFG spots (~70% of total cellular TFG) colocalized with Sec31, although a considerable number of TFG spots, which were typically dimmer, did not localize at ERES (negative for Sec31). Upon incubation of HSC3 cells with 10 ng/ml EGF-Rh, the extent of TFG:Sec31 colocalization showed a slight transient decline (Figure 5G), whereas about 10% of total cellular TFG was colocalized with EGF-Rh containing endosomes (Figures 5B and H). The extent of TFG colocalization with EGF-Rh may have been however underestimated because of poor fixability of EGF-Rh (lacking any free amino groups) and its loss during cell permeabilization. Visual analysis of TFG and EGF-Rh fluorescence suggested that EGF-Rh is mostly co-localized with Sec31-negative, dim TFG puncta (Figure 5B). Nevertheless, a small fraction of EGF-Rh endosomes was seen colocalized with or juxtaposed to bright Sec31-positive TFG puncta (Figure 5B). In summary, the data in Figures 4 and 5A-B, G-H show that whereas, consistently with previous studies, the bulk of TFG is located at ERES, a smaller but substantial pool of TFG is either associated with EGFR-containing endosomal compartments or located in close proximity to endosome membranes.

**Figure 5.**
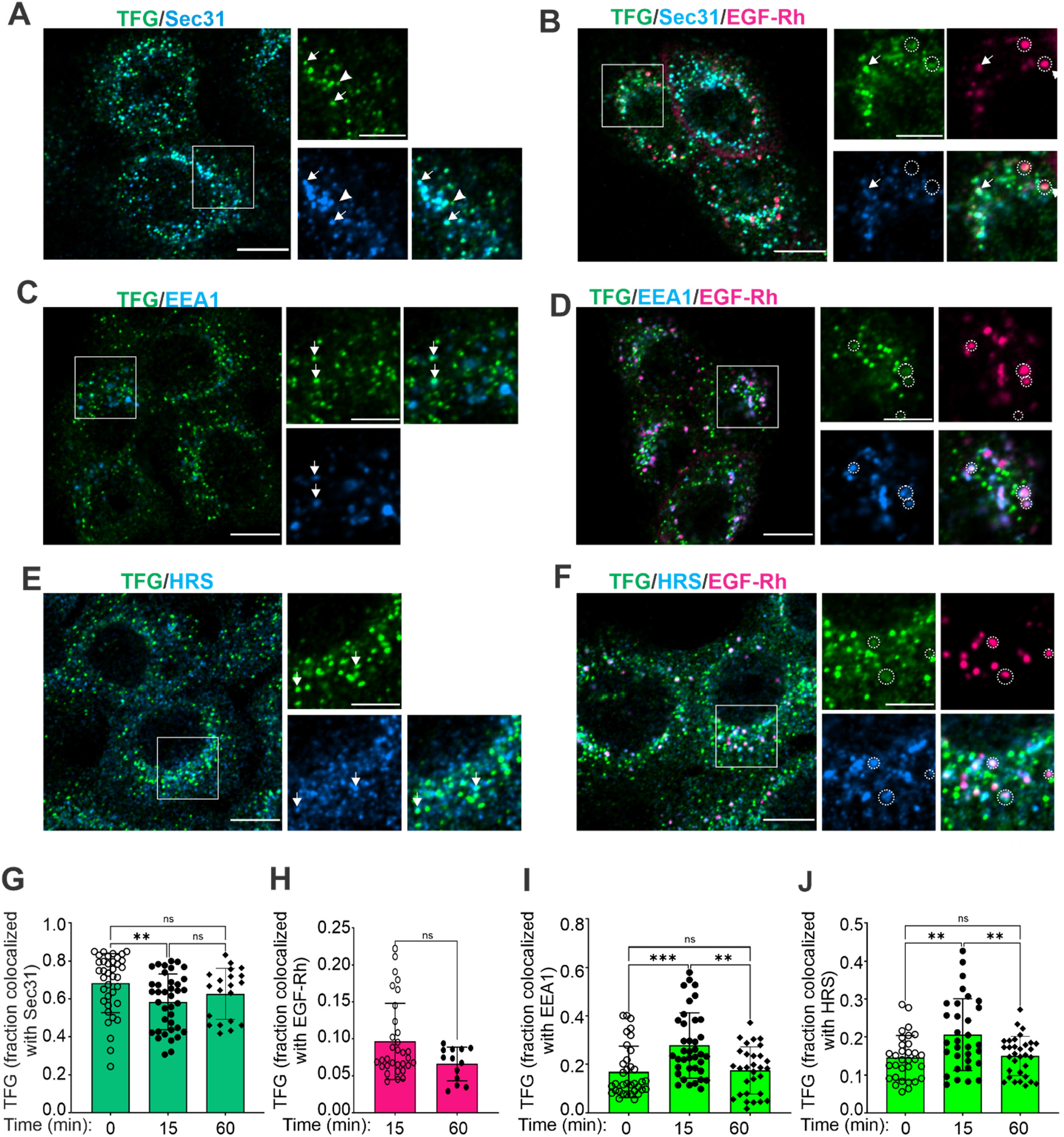
Colocalization of TFG with ERES, EGF-Rh and endosomal markers. HCS3 cells were untreated or treated with 10 ng/ml EGF-Rh for 15 min or 60 min (images are not shown) and fixed. (**A**-**B**) Staining with TFG and Sec31 antibodies. 3D confocal images were acquired through 640 nm (*cyan*, Sec31), 561 nm (*magenta*, EGF-Rh) and 488 nm channels (*green*, TFG). Arrows point at examples of TFG localization with Sec31, and arrowheads point at TFG puncta that is not co-localized with Sec31. Circles indicate examples of colocalization of TFG with EGF-Rh outside of ERES. (**C-D**) Staining with TFG and EEA1 antibodies. 3D confocal images were acquired through 640 nm (*cyan*, EEA.1), 561 nm (*magenta*, EGF-Rh) and 488 nm channels (*green*, TFG). Arrows point at examples of TFG localization with EEA1. Circles indicate colocalization of TFG, EEA1 and EGF-Rh. (**E-F**) Stained with TFG and HRS antibodies. 3D confocal images were acquired through 640 nm (*cyan*, HRS), 561 nm (*magenta*, EGF-Rh) and 488 nm channels (*green*, TFG). Arrows point at examples of TFG localization with HRS. Circles indicate colocalization of TFG, HRS and EGF-Rh. Insets in ***A-F*** represent enlargements of the regions indicated by rectangles. All images are single sections in the middle of the cell of representative 3D images. Scale bars, 10 μm in full images and 5 μm in insets. (**G**) Quantification of TFG-Sec31 co-localization from images exemplified in ***A-B***. Bar graph represents mean values with SDs (n 20-37 FOVs from 3-6 independent experiments). Kruskal-Wallis test. Dunn’s multiple comparisons test. (**H**) Quantification of the fraction of TFG colocalized with EGF-Rh from images exemplified in ***A-B***. Bar graph represents mean values with SDs (n of 13-32 FOVs from 2-5 independent experiments). Mann-Whitney test. (**I**) Quantification of TFG-EEA1 co-localization from images exemplified in ***C-D***. Bar graph represents mean values with SDs (n of 30-38 FOVs from 4-6 independent experiments). Kruskal-Wallis test. Dunn’s multiple comparisons test. (**J**) Quantification of TFG-HRS co-localization from images exemplified in ***E-F***. Bar graph represents mean values with SDs (n of 31-32 FOVS from 4 independent experiments). One-way ANOVA. Tukey’s multiple comparison test.

### Fraction of TFG in early/sorting endosomes is increased by EGF

To address whether TFG targeting to early/sorting endosomes is EGF-dependent, untreated and EGF-Rh-treated HSC3 cells were co-stained with the TFG and EEA1, HRS or Rab5 antibodies. TFG was partially colocalized with EAA1-positive endosomes in both untreated and stimulated cells (Figure 5C-F). Quantifications revealed an ~1.5-fold increase in the extent of TFG colocalization with EEA1 after 15 min of EGF-Rh stimulation followed by the decrease in this colocalization after 60 min down to the extent measured in untreated cells (Figure 5I). Similar trend of increased colocalization of TFG with HRS (Figure 5J) in EGF-Rh-stimulated cells was also observed. The fraction of TFG colocalized with Rab5 was smaller (<5%) than that colocalized with EEA1 (20-30%) or HRS (15-20%), however, EGF-dependent increase of TFG co-localization with Rab5 was also observed (Figure S2A-C). In all experiments, internalized EGF-Rh was highly co-localized with markers of early/sorting endosomes. On the other hand, accumulation of ligand-free EGFR in endosomes triggered by activation of stress-induced p38-MAPK with 100 nM Anisomycin in HSC3 cells did not increase colocalization of TFG and EEA1 (Figure S2 D-F). Collectively, Figures 5 and S2 demonstrate that TFG is constitutively associated with early/sorting endosomes and that ligand activation of EGFR enhances this association. It should be noted that visual inspection of individual examples of TFG localization in endosomes shows that shapes of the puncta labeled with EGF-Rh or endosomal markers, and shapes of an overlapping TFG-labeled puncta are often not the same. This may be due to localization of TFG, EGF-Rh and endosomal markers in different subdomains of endosomes, such as for example, intralumenal vesicles (possible location of EGF-Rh) and so called Rab5-EEA1 and HRS subdomains (Raiborg et al., 2006). Alternatively, a fraction of TFG may be not directly associated with endosomes but located in close proximity to endosomal membranes (<20 nm), which would result in an overlap of corresponding fluorescence signals that is not resolved by confocal microscopy and therefore scored as co-localization.

### TFG depletion accelerates EGF-induced EGFR degradation and causes enlargement of endosomes

To begin an investigation of the TFG function in endosomes and EGFR endocytic trafficking, we examined effects of siRNA knockdown of TFG on EGF-induced EGFR degradation. Figures 6A and C show that TFG depletion with validated siRNA duplex (Johnson et al., 2015) was highly efficient (>90%). Surprisingly, TFG knockdown increased rates of ligand-induced EGFR degradation in both HSC3 and HeLa/FAP-EGFR cells (Figure 6A-D). Turnover of ligand-activated EGFR in HeLa/FAP-EGFR cells was substantially faster than in HSC3 cells (mean t1/2 values of 0.65 hrs versus 8.4 hrs, respectively), and therefore, additional acceleration of such fast degradation process by TFG knockdown was less pronounced in HeLa/FAP-EGFR than in HSC3 cells. In contrast, TFG depletion did not alter constitutive EGFR degradation in both cell lines (Figure S3A-D). These data suggest that TFG depletion does not have a pleiotropic effect on endocytic and anterograde trafficking, but that its degradation-accelerating effect is specific for endocytic cargo such as ligand-activated endosomal EGFR.

**Figure 6.**
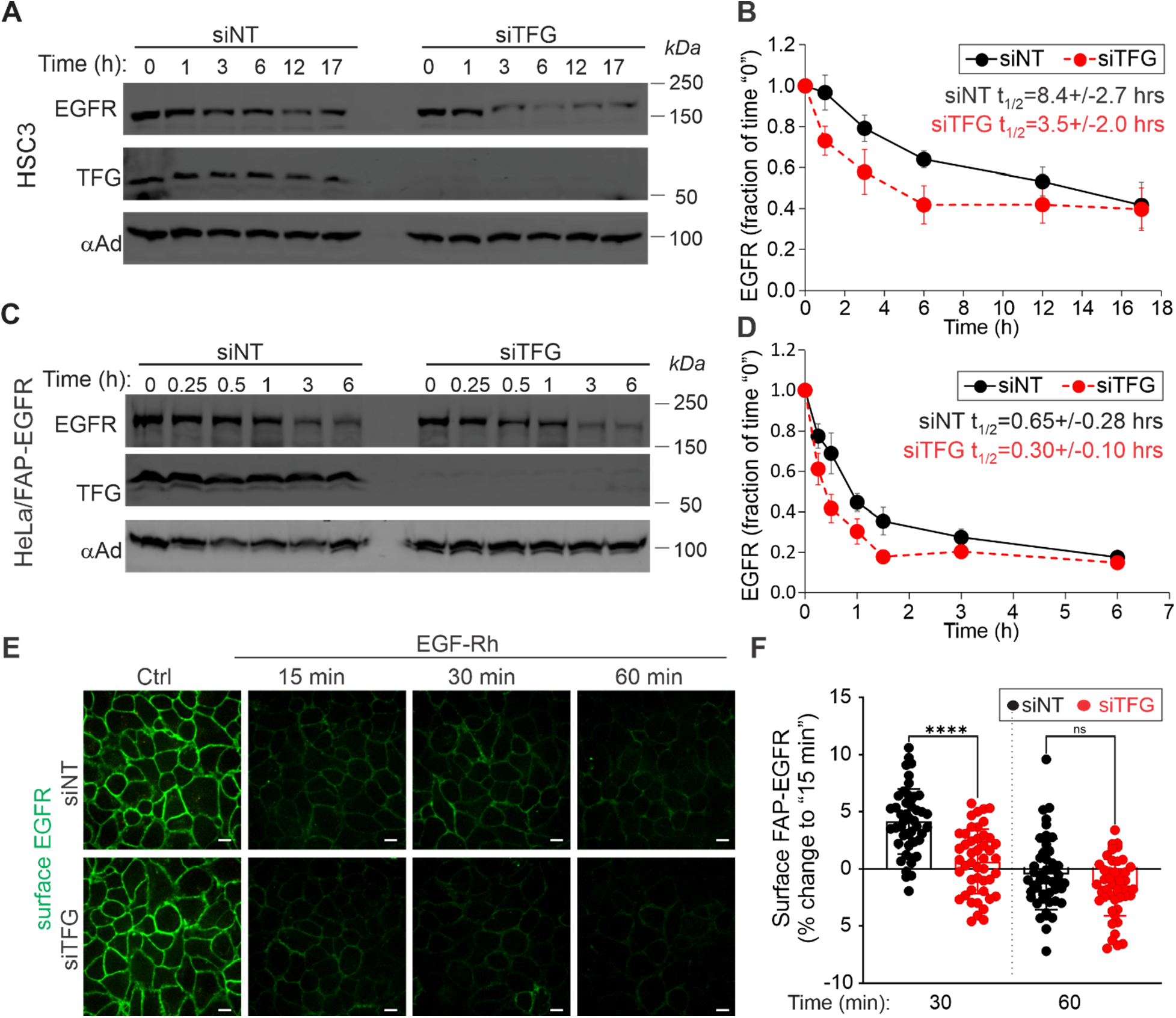
TFG depletion by siRNA accelerates EGF-induced EGFR degradation. (**A**-**B**) HSC3 and (**C**-**D**) HeLa/FAP-EGFR cells were transfected with non-targeting (siNT) or TFG (siTFG) siRNAs. After 3 days, cells were preincubated with 20 μg/ml cycloheximide for 1 h and incubated with 10 ng/ml EGF for indicated times in the presence of cycloheximide. Cells were then lysed, and the lysates were probed by Western blotting with antibodies to EGFR, TFG and α-adaptin (*αAd*, loading control). In ***C*** and ***D***, the amount of EGFR was normalized by the amount of α-adaptin, expressed as fraction of the normalized value at time “0”, plotted against time and presented as mean values with SEMs. Half-life time of EGFR degradation (t1/2) was calculated using one phase decay least squares fit and presented as mean values with SDs. (n is 4 and 9 independent experiments in HSC3 and HeLa/FAP-EGFR cells, respectively). (**E-F**) HeLa/FAP-EGFR cells transfected with siNT or siTFG were incubated with 10 ng/ml EGF-Rh for indicated times. Internalization was stopped by placing the cells on ice, and surface FAP-EGFR was labeled with MG-B-Tau. Cells were fixed and imaged through the 640 nm channel (*green*, surface EGFR). Representative sections in the middle of the cells are shown. Intensity scales are identical on all images. Scale bars, 10 μm. In ***F***, fluorescence intensities of MG-B-Tau per FOV were quantified. The mean value obtained at 15 min was subtracted from each FOV value at indicated time points and the resulting difference normalized by the mean value from untreated cells to obtain the percentage change of the fluorescence intensity. Bar graph shows mean with SDs (n of 46-47 FOVs from 5 independent experiments). Kruskal-Wallis test. Dunn’s multiple comparison test.

The effect of TFG knockdown on ligand-induced EGFR degradation could be explained by increased lysosomal targeting and degradation and/or decreased recycling of EGFR from endosomes to the plasma membrane. To test whether TFG affects EGFR recycling, we used HeLa/FAP-EGFR cells, which were untreated or treated with 10 ng/ml EGF-Rh for 15, 30 or 60 min before labeling FAP-EGFR located at the cell surface with the membrane impermeable fluorogen MG-B-Tau at 4°C (Perez Verdaguer et al., 2021). Incubation of cells with EGF-Rh for 15 min reduced the amount of FAP-EGFR at the cell surface by ~40% in both siNT vs siTFG transfected cells (Figure 6E). This down-regulation was followed by a small transient recovery in the next 15 min (30 min time-point) in control cells but not in TFG-depleted cells (Figure 6E-F). Such transient increase in the cell-surface EGFR level was mediated by endosomal recycling rather than insertion of newly synthesized EGFRs, as the anterograde traffic is too slow to significantly contribute to the increase in cell-surface EGFR within 15 min (Scharaw et al., 2016). Interestingly, when HeLa/FAP-EGFR cells were stimulated with 10 ng/ml TNFα, which transiently activates p38-mediated internalization of ligand-free EGFR, TFG depletion did not alter EGFR recycling (Figure S3E-F). The fluorescence intensity of internalized EGF-Rh after 15 min incubation of control and TFG-depleted cells was similar (Figure S3G), indicating that increased EGF-induced EGFR degradation in TFG-depleted cells was not due to increased internalization. Therefore, the data in Figures 6 and S3 imply that accelerated degradation of EGFR in TFG-depleted cells is at least in part due to an impaired recycling and suggest that TFG may regulate endosomal sorting of EGFR.

To further explore TFG role in endosomal sorting, we tested whether TFG knockdown affects endosomal labeling of EEA1 and HRS. Surprisingly, early/sorting endosomes appeared brighter and apparently larger in TFG-depleted than in control cells (Figure 7). Differences in the endosome appearance were especially revealing in images depicting cells depleted and not fully depleted of TFG by siRNA next to each other (Figure 7B and D). Quantifications showed that TFG knockdown increased the apparent volume of EEA1 and HRS containing early/sorting endosomes (Figures 7C-D and G-H). To further substantiate these observations, we performed analysis of endosomal compartments using correlated light and electron microscopy (CLEM). Control and TFG-depleted HSC3 cells were incubated with 200 nM ferritin for 30 min to label endosomes. High-contrast monochrome light microscopy (LM) images were acquired from ultrathin sections, and electron microscopy (EM) images were acquired from multiple cell regions chosen on LM images to verify that ferritin positive compartments display the morphology of early endosomes and multivesicular (sorting) endosomes (Supplemental Figure S4A-C). AI-based quantification of LM images revealed that TFG-depleted cells tend to have increased number of endosomes and larger endosomes when compared to cells transfected with non-targeting siRNA (Figure S4D-E). Together, fluorescence microscopy and CLEM suggest a role of TFG as a modulator of a general organization of the endolysosomal system. Considering that TFG appears to have a “pro-recycling” function in endosomal sorting of cargo like EGFR (Figure 6), it is possible that impaired removal of recyclable components from sorting endosomes results in “build-up” of these components in endosomes, thus leading to enlargement and accumulation of endosomes in the cell.

**Figure 7.**
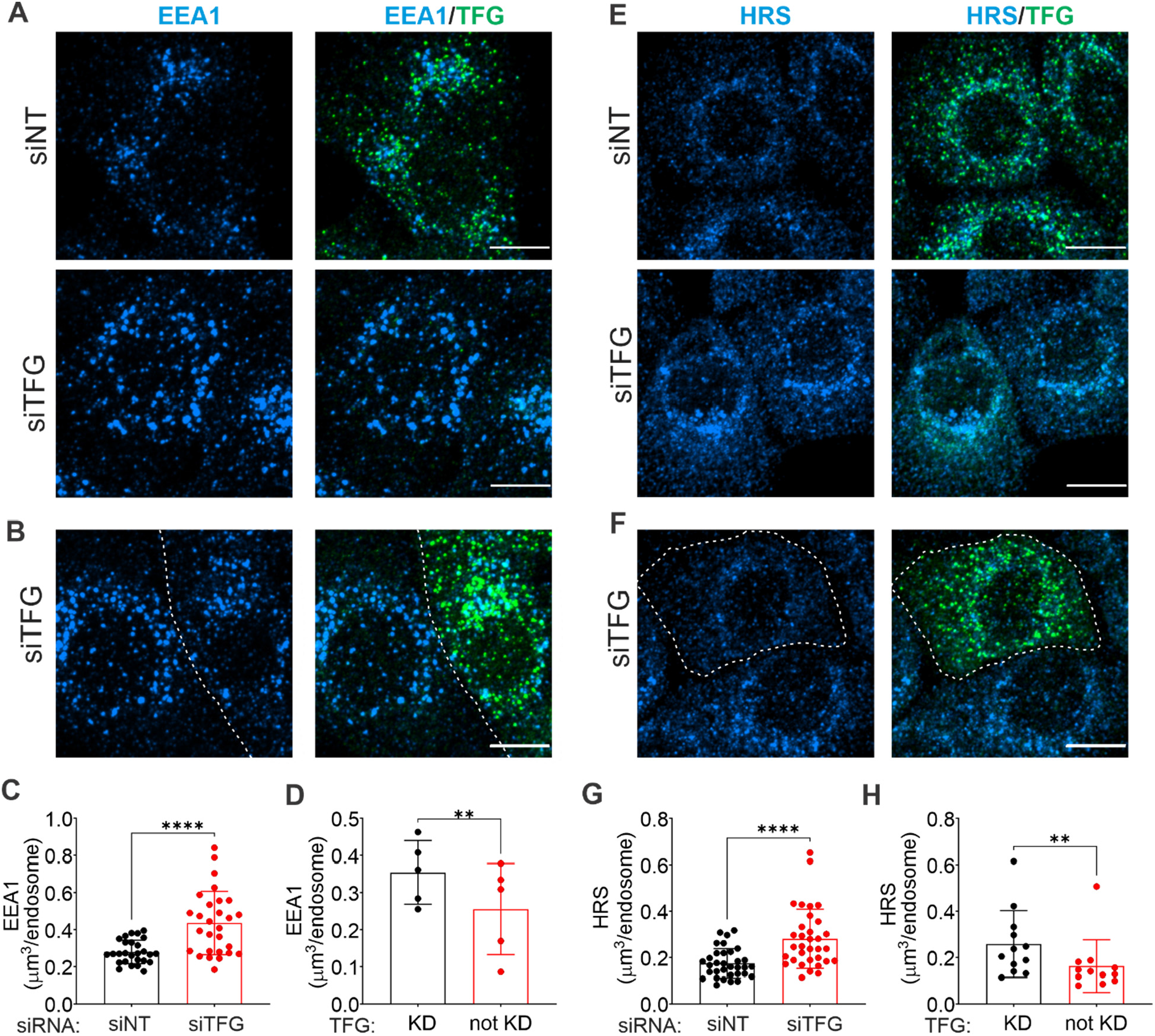
TFG siRNA depletion causes apparent enlargement of early/sorting endosomes. HSC3 cells transfected with non-targeting (siNT) or TFG siRNAs (siTFG) were fixed and co-immunolabeled with TFG and EEA1 (**A**-**D**) or HRS antibodies (**E-H**). 3D confocal images were acquired through 640 nm (*cyan*, EEA1 and HRS) and 488 nm channels (*green*, TFG). Representative maximum intensity projections of 3D images are shown. Scale bars, 10 μm. In **C-D** and **G-H**, the apparent volume of EEA1 and HRS labeled endosomes was calculated in images exemplified in **A-B** and **E-F**. Bar graphs show mean values with SDs (**C**: n of 27-28 FOVs from 4 independent experiments. Mann-Whitney test; **D**: n of 5 FOVs from 2 independent experiments. Paired t-test; **G**: n of 33 FOVs from 4 independent experiments. Mann-Whitney test; **H:** n of 12 from 3 independent experiments. Wilcoxon matched-pairs test).

After we began initial studies of TFG function in EGFR-containing endosomes, it was reported that TFG is involved in Wnt signaling and colocalizes with HRS (Colozza et al., 2020). Most recently, a role of TFG in ER-associated autophagy was proposed, further supporting potential involvement of TFG in the crosstalk of the endolysosomal system and ER (Carinci et al., 2021). However, another study suggested that ERES-ERGIC associated autophagy does not require TFG (Li et al., 2021). We speculate that the TFG function in the endosomal system, possibly promoting recycling, may be somewhat similar to the proposed role of TFG in COPII vesicle uncoating and spatially organizing the ERES-ERGIC trafficking “hub”. In sorting endosomes, TFG may organize and facilitate exit of recycling carriers. Undoubtedly, while more studies beyond our initial analysis are needed to precisely define the specific function of TFG in endosomes, identification of TFG as a new regulator of endosomal sorting illustrates the power of APEX2-based time-resolved analysis of the proximity proteomes of endocytosed receptors for discovering new mechanisms of their signaling and endocytic trafficking.

## Materials and Methods

### Reagents

Recombinant human EGF was purchased from BD Biosciences (San Jose, CA). EGF-Rh was from Molecular Proves (Invitrogen, Carlsbad, CA). TNFα was from Miltenyi Biotec (Germany). Anisomycin (stored as 1 mM stock solution in DMSO at 4°C), was from Sigma-Aldrich (Saint Louis, MO). Rabbit polyclonal antibody to EGFR (AB_2246311) was from Cell Signaling Technology (Danvers, MA). Mouse monoclonal antibody to EGFR (AB_10978829) was from American Type Culture Collection. Rabbit polyclonal antibody to Hrs (AB_1124436) was from Santa Cruz Biotechnology (Dallas, TX). Monoclonal IgG2a antibody to α-adaptin (α subunit of AP2) (AB_2056321) was from ThermoFisher (Waltham, MA). Mouse monoclonal antibodies to EEA1 (AB_397829), Sec31 (AB_399717) and Rab5 (AB_398047) were from BD Transduction (Franklin Lakes, NJ). Rabbit polyclonal antibody to TFG (AB_2664282) was from Invitrogen. Secondary anti-mouse and anti-rabbit IRDye antibodies were from Li-COR (Lincoln, NE). Secondary anti-mouse and anti-rabbit conjugated to Alexa488 or Alexa640 were from Jackson immune (West Grove, PA). MG-B-Tau was provided by Dr. M. Bruchez (Carnegie Mellon University). Ferritin (F4503) was from Sigma-Aldrich.

### Expression plasmids

We created plasmids to express APEX2 tagged EGFR in HCT116 and HEK293T cells. Gateway entry clones pDONR223-EGFR-WT (#81926) were purchased from Addgene. pLEX-305 (#41390) lentiviral backbone vector was engineered to substitute the hPKG (phosphoglycerate kinase) promoter with the CMV (human cytomegalovirus) promoter plus APEX2 tag in the C-terminus of the expression gene. pDONR223_EGFR_WT (#81926) were transferred to pLEX305_Cterm_APEX2 destination vector using Gateway LR Clonase II enzyme mix (Thermo Fisher) based on the provided protocol. Gateway cloning product was transformed into One Shot™ BL21(DE3) chemically competent *E.coli* (Thermo Fisher). GFP-TFG was provided by Dr. Hideki Shibata (Nagoya University, Chikusa-ku, Nagoya, Japan)

### Cell culture

HEK293T, HCT116 cells (ATCC) and HeLa/FAP-EGFR (Larsen et al., 2019) were maintained with DMEM (Gibco) supplemented with 10% FBS (v/v) and 50 U/ml penicillin and 50 U/ml streptomycin (Gibco). HSC3 cells were maintained in DMEM with 5% FBS. For all experiments cells were serum-starved 16-24 hrs before experimental treatments: HEK293T and HSCT116 cells culture medium was replaced by DMEM supplemented with 1% FBS and Penicillin-Streptomycin; for HeLa/FAP-EGFR and HSC3 cell culture medium was replaced by DMEM. All experimental treatments were carried out in DMEM supplemented with 0.1% BSA at 37°C.

### DNA plasmid transfections, generation of stable cell lines and RNA interference

HeLa/FAP-EGFR cells were transfected with the GFP-TFG plasmid using Lipofectamine 2000 (ThermoFisher Scientific) following manufacturer’s instructions and used for experiments 24 hrs later. For generation of EGFR-APEX2 stable cell lines, one day before transfection, we seeded HEK293T cells at 70% to 90% confluence. 5 μg validated clones were combined with viral helper constructs (VSVG, TAT1B, MGPM2, CMV-Rev1B) at a ratio 4:1 and diluted in 500 μl Opti-MEM with 30 μL P3000 Reagent. Diluted DNA was added to 500 μL Opti-MEM with 22 μl Lipofectamine 3000 reagent. The mixture was incubated at room temperature for 15 min and then added to cells. The cells were grown for three days for virus production. Virus was collected and filtered using Millex-GP Syringe Filter Unit, 0.45 um (EMD Millipore). Various amount of collected virus were added to HEK293T or HCT116 cells. After 48 hrs, 2 μg/ml and 0.5 μg/ml puromycin were added for selection of single clones of HEK293T and HCT116 cells expressing EGFR-APEX2, respectively.

Non-targeting siRNA was purchased from Qiagen (Venlo, Netherlands) and TFG targeting siRNA duplexes (individual siGENOME grade) purchased from Dharmacon (Lafayette, CO): Sense sequence:

5’-CCAAAAGACUCCAGUACUAUU

Antisense sequence: 5’-UAGUACUGGAGUCUUUUGGUU

HSC3 or HeLa/FAP-EGFR cells were reverse transfected in 6-well dishes with 50 nM siRNA and 3.5 μl DharmaFECT-1 in 1 ml of complete media without antibiotics. 2 days later, cells were split and plated in coverslips or other dishes (for western blotting or EGFR surface recovery analysis). 6 hrs after, media was changed to serum-free DMEM, and the cells were used for experiments the next day (day 3 after siRNA transfection).

### Western blotting

Cells were lysed and processed for Western blotting as described in (Larsen et al., 2019) and (Paek et al., 2017), in Sorkin and Gygi laboratories, respectively. Quantifications of band intensities were performed using a Li-COR software.

### Proximity-based labeling

After a 24 hrs starvation, biotinyl tyramide (BT; Toronto Research Chemicals) stock was added directly into starvation media to a final concentration of 500 μM. Cells were incubated in labeling medium for 1 h. EGF was added to treat the cells for various times. An equal volume of PBS buffer (vehicle) was added to control cells. Hydrogen peroxide was freshly diluted from a 2M H_2_O_2_ stock and added into the medium to a final concentration of 1 mM 59 min after the start of incubation with BT to initialize the labeling reaction. After 1 min of H_2_O_2_ treatment, cells were washed with ice-cold quenching solution (PBS supplemented with 10 mM sodium ascorbate, 5 mM trolox, and 10 mM sodium azide) thrice and quenching solution with 5 mM EDTA once. 1 ml ice-cold lysis buffer (2 M sodium hydroxide with 7.5% 2-mercaptoethanol in Milli-Q water) was added directly to the plate to harvest the cells. Cell lysates were stored at −80°C until the next step.

### Streptavidin pull-down of biotinylated proteins

Cell lysates were syringe lysed to fragment DNA (20x with 21gauge needle), 0.7 ml ice-cold trichloroacetic acid (TCA) was added, and the samples were incubated on ice for 15 min. Proteins were precipitated by centrifugation at 21,130 × g at 4°C for 10 min. Pellets were washed with −20 °C cold acetone, vortexed, and centrifuged at 21,130 × g at 4 °C for 10 min. This step was repeated with ice-cold methanol. Methanol was aspirated and 0.8 ml resuspension buffer (8M urea, 100 mM sodium phosphate pH 8, 100 mM NH_4_HCO_3_, 1% SDS (v/v)), were added to the pellets. The pellets were vortexed and suspended in the lysis buffer. A BCA assay determined the protein concentration and an equal amount (2.5-3mg) of protein was used for each sample for the following steps. Samples were reduced with 5 mM TCEP, alkylated with 10 mM iodoacetamide, and quenched with 15 mM DTT. 50 mM ammonium bicarbonate was added to each sample to reach final concentrations of 4 M urea and 0.5% (vol/vol) of SDS. A 75 μl suspension equivalent per sample of streptavidin magnetic beads (Thermo Fisher Scientific, stock is at 10mg/ml, 100 μl can bind 55 μg) were washed twice with 4M urea, 0.5% SDS (vol/vol), 100 mM sodium phosphate pH 8, 75 μl of beads were added to each sample, and tubes were rotated for 3 hrs at RT. Following the streptavidin pull-down, magnetic beads were washed thrice with 4 M urea, 0.5% SDS (vol/vol), 100 mM sodium phosphate pH 8, once with the 4 M urea, 100 mM sodium phosphate pH 8, and once with 200 mM EPPS pH 8.5 sequentially to remove the SDS in the samples.

### On-beads digestion and TMT labeling

Washed beads were re-suspended in 100 μl EPPS pH 8.5. 1μl of LysC stock solution (2mg/ml, Wako) was added into each sample. Samples were vortexed briefly and incubated at 37°C for 3 hrs with shaking. Trypsin stock (Promega #V51113) 1:200 (vol/vol) was then added for further digestion overnight at 37°C with shaking. Magnetic beads were removed from the samples. 30 μl acetonitrile were added into each sample. 100 μg of each TMT 10plex (or TMT 9/11plex) reagent were used for labeling. After 1 h labeling, 2 μl of each sample were combined, desalted, and analyzed using mass spectrometry. Total intensities were calculated in each channel to determine normalization factors. After quenching using 0.3% hydroxylamine, samples were combined at a 1:1 ratio of peptides based on normalization factors, dried, and fractionated using High pH Reversed-Phase Peptide Fractionation kit (Pierce). Six fractions were dried, desalted, and analyzed by liquid chromatography-tandem mass spectrometry (LC-MS/MS).

### LC-MS/MS

The data acquisition method was as described previously (Navarrete-Perea et al., 2018). Briefly, mass spectrometric data were collected on an Orbitrap Fusion Lumos mass spectrometer coupled to a Proxeon NanoLC-1200 UHPLC. The 100 μm capillary column was packed in-house with 35 cm of Accucore 50 resin (2.6 μm, 150Å; ThermoFisher Scientific). The mobile phase was 5% acetonitrile, 0.125% formic acid (A) and 95% acetonitrile, 0.125% formic acid (B). The data were collected using a DDA-SPS-MS3 method. Each fraction was eluted across a 120 min method with a gradient from 6% to 30% B. Peptides were ionized with a spray voltage of 2,600 kV. The instrument method included Orbitrap MS1 scans (resolution of 120K; mass range 350-1400 m/z; automatic gain control (AGC) target 5×10^5^, max injection time of 50 ms) and ion trap MS2 scans (isolation window: 0.5, CID collision energy of 35%; AGC target 1×10^4^; rapid scan mode; max injection time of 60 ms). MS3 precursors were fragmented by HCD and analyzed using the Orbitrap (NCE 65%, AGC 3 x10^5^, maximum injection time 150 ms, resolution was 50,000 at 400 Th). The mass spectrometry proteomics data have been deposited to the ProteomeXchange Consortium via the PRIDE (Perez-Riverol et al., 2019) partner repository with the dataset identifier XXX. Data are available via ProteomeXchange with identifier PXD030072.

### Data analysis

Mass spectra were processed using a SEQUEST-based software pipeline and searched against the human UniProt database (Downloaded in 2014-02-14). Searches were performed using a 50-ppm precursor ion tolerance and 0.9 Da ion tolerance. TMT tags on lysine residues and peptide N termini (+229.163 Da) and carbamidomethylation of cysteine residues (+57.021 Da) were set as static modifications. Oxidation of methionine residues (+15.995 Da) was set as a variable modification. Peptide-spectrum matches (PSMs) were identified, quantified and filtered to a 1% peptide false discovery rate (FDR). In APEX protein quantitation, peptides were collapsed to a final protein-level FDR of 1%. Proteins were quantified by summing reporter ion counts across all matching PSMs. Briefly, a 0.003 Da (3 millidalton) window around the theoretical m/z of each reporter ion was scanned and the maximum intensity nearest the theoretical m/z was used. For each protein, signal-to-noise (S:N) measurements of the peptides were summed, and these values were normalized to the EGFR level in each channel. Proteins with summed signal-to-noise greater than 100 were used for further analysis.

### Immunofluorescence microscopy

Cells were grown on glass coverslips, and after indicated treatments, fixed in freshly prepared 4% paraformaldehyde (PFA) for 15 min, washed with Ca^2+^, Mg^2+^-free PBS (CMF-PBS), permeabilized in 0.1% Triton X-100/CMF-PBS for 15 min and blocked for 30 min in 0.1% BSA in CMF-PBS. Cells were then incubated for 1h with primary and secondary antibodies in blocking buffer. Samples were mounted in Mowiol (Calbiochem). All procedures with fixed cells were carried out at RT.

### Detection and measurement of cell-surface FAP-EGFR

HeLa/FAP-EGFR cells were grown in glass coverslips. Cells were treated with 10ng/ml EGF-Rh or 10ng/ml TNFα at 37°C for indicated times, washed with cold CMF-PBS and incubated with 100 nM MG-B-Tau in CMF-PBS for 10 min on ice. Finally, cells were washed in CMF-PBS and fixed with 4% freshy prepared PFA for 15 min at RT, washed in CMF-PBS and imaged immediately.

### Spinning disk confocal microscopy

Images were acquired with a spinning-disk Marianas system based on a Zeiss Axio Observer Z1 inverted fluorescence microscope equipped with 63x Plan Apo PH NA 1.4 oil immersion objective, piezo stage controller, spherical aberration correction module, temperature- and CO_2_-controlled chamber, all controlled by Slidebook6 software (Intelligent Imaging Innovation, Denver, CO) as described previously (Larsen et al., 2019; Perez Verdaguer et al., 2019). For immunofluorescence imaging, Hamamatsu Flash4 camera was used to obtain z-stack of 15 x-y confocal images acquired at 250nm z-steps. For surface FAP-EGFR and internalized EGF-Rh measurements, an Evolve EM-CCD camera was used to obtain single image in the middle of the cells. Image acquisition settings were identical for all variants in each experiment. Typically, 6-10 fields of view (FOV) including several cells were imaged per condition in each experiment.

### Image analysis

To quantify the amount of TFG localized at ERES or in endosomes, a colocalization analysis was performed using Slidebook6. A segment mask was generated to select ERES (Sec31), early/sorting endosomes (EEA1, HRS, Rab5) or EGF-Rh vesicles detected through the 640 nm (Alexa647) or 561 nm (rhodamine) channels (Target Compartment mask) in background subtracted images. Another segment mask was generated to select TFG-positive voxels in the 488 nm channel (TFG mask). For both masks identical threshold parameters were used for all experimental variants. Objects smaller than 6 voxels were eliminated. A Colocalization mask was then generated to select voxels overlapping in the Target Compartment and TFG masks. The sum fluorescence intensity through the 488 nm channel in the Colocalization mask was divided by the sum fluorescence intensity of the TFG mask in each FOV.

To quantify the apparent volume of endosomes, the total volume of Target Compartment mask was divided by the number of objects in the mask. To measure the amount of cells-surface FAP-EGFR, the sum intensity of the 640 nm channel (MG-B-Tau) per FOV was obtained. The mean value of MG-B-Tau fluorescence intensity at 15 min was subtracted from each value at 30 min and 60 min of EGF-Rh stimulation and then normalized by the mean value of this intensity obtained in untreated cells to obtain the recovery fraction. To measure EGF-Rh endocytosis, the sum intensity of 561 nm channel (EGF-Rh) per FOV was obtained and normalized by the mean value of 561 intensity in siNT cells.

### CLEM

Cell incubated with ferritin were fixed for 30 min in 2.5% glutaraldehyde (Taab) in PBS, and washed 3x in PBS before postfixing in 1% osmium tetroxide for 1 hr. Following dehydration through a graded series of alcohols, epoxy resin Beem capsules full of resin were inverted over random areas of the culture dish and the resin cured for 24 hrs. The resin filled Beem capsules were snapped of the dish, trimmed to a trapezoid and sections cut with a Leica UltraCut R (Leica Microsystems Mannheim Germany) microtome. Ultrathin (60 nm) sections were mounted on Formvar coated 3 slot copper grids and stained with uranyl acetate and lead citrate (Watkins and Cullen, 1986), serial 200 nm thick sections were mounted on glass slides and stained with 1% Toluidine blue for 5 min.

The high contrast intracellular nature of ferritin positive structures within semi-thin sections of EM blocks viewed by light microscopy allows the use of neural network training of segmentation of the ferritin positive structures within sections. Monochromatic brightfield image montages (3×3 images) were collected using a carefully aligned Nikon Ti microscope (Nikon instruments, Tokyo Japan) with a 100x 1.49 NA TIRF objective and a 1.5x intermediate magnifying lens collected to a Teledyne Photometrics 95B camera (Photometrics,Tucson Arizona) with 1800×1800 pixels and a 25 x25 mm chip sensor (11 μm pixels). Final calibrated pixel size is 0.07 μm on each axis. To generate correlative images between light and EM modalities, low magnification images (20x 0.75NA) of the entire section were collected and used to find reference areas in the serial EM sections, such that the identity of ferritin positive structures could be confirmed. EM images were collected using a JEOL 1200 Flash (Peabody Ma) electron microscope.

To quantify the number and size of ferritin positive structures (endosomes) in each cell and treatment condition, we used NIS Elements and the AI segmentation tool. A ground truth data set was generated by manually highlighting and defining structures in the monochrome LM images. A second training set was used to segment out the cell boundaries such that following training we are able to collect the number of ferritin positive structures and size (area) of individual ferritin-positive structures in each cell. Five 3×3 montages were collected for each condition and the entire montages were passed through the trained segmentation algorithm. In total around 700 cells per condition were analyzed. Only cells within the field of view and with nuclei were included in the analysis. To show the distribution of the number of endosomes per cell within cell population, cells were classified in arbitrary categories according to their endosome number. To show the distribution of endosome size within cell population, averaged area of individual endosomes in each cell was quantified, and the cells were classified in arbitrary categories of this averaged area of endosomes.

### Statistics

Statistical analyses of fluorescence microscopy and blotting experiments were performed using GraphPad Prism software (GraphPad). We tested our datasets for normal distribution with Shapiro-Wilk test and chose an appropriate test accordingly. We used unpaired Student’s t-test or Mann-Whitney test for comparisons of two samples with equal or unequal variance respectively. In Figure 7D and H paired Student’s t-test or Wilcoxon test were used. For comparison of multiple groups, a one-way ANOVA or Kruskal-Wallis test respectively followed by Tukey’s or Dunn’s post hoc tests were used. All error bars denote mean+/- SD unless otherwise indicated in figure legend. *p<0.05, **p<0.01, ***p<0.001. The number of individual experiments analyzed are indicated in the figure legends.

## Acknowledgements

We appreciate gifts of reagents by Drs. Bruchez and Shibata. We are grateful to Ms. M. Sullivan for technical help with EM. This work was supported by NIH grants GM124186 and CA089151.

