## Supplemental Figures for "Time-resolved proximity labeling of protein networks associated with ligand-activated EGFR"

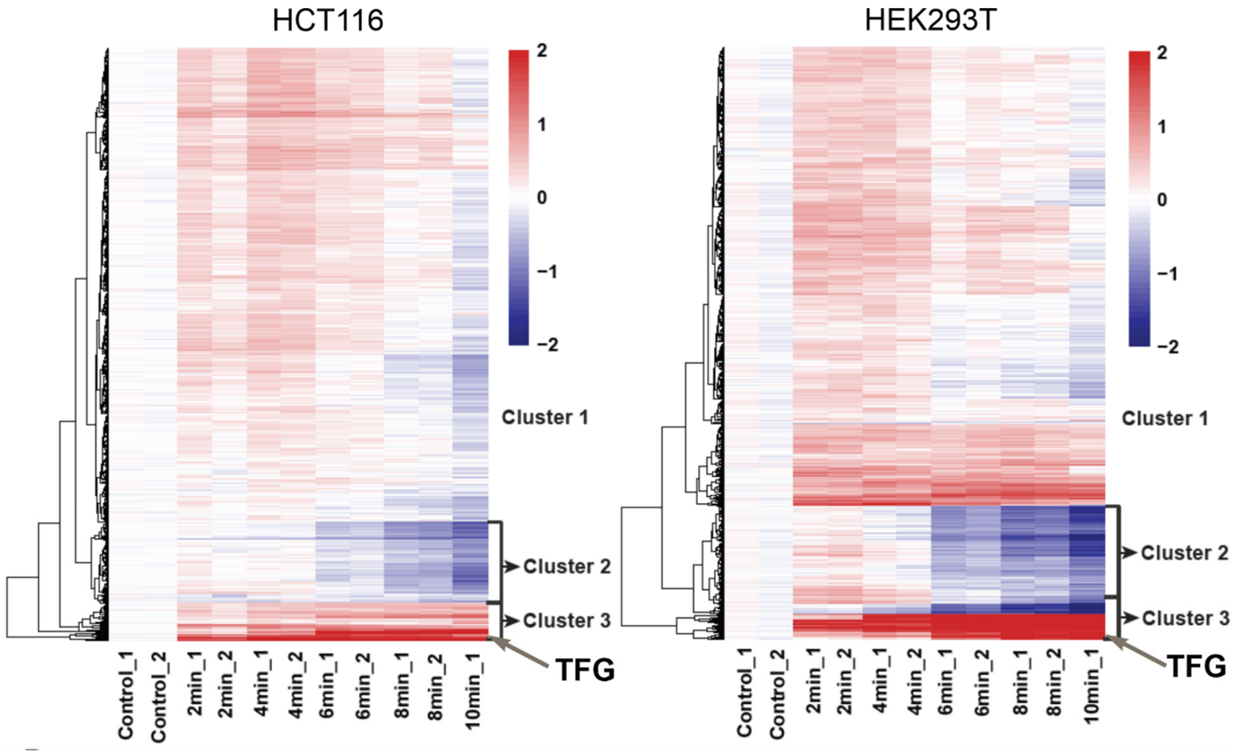


**Figure S1. Heatmaps of proteins quantified in the EGFR-APEX2 experiment in Figure 4A.** HCT116 cells (**A**) and HEK293T cells (**B**). Cluster analysis using K-means.

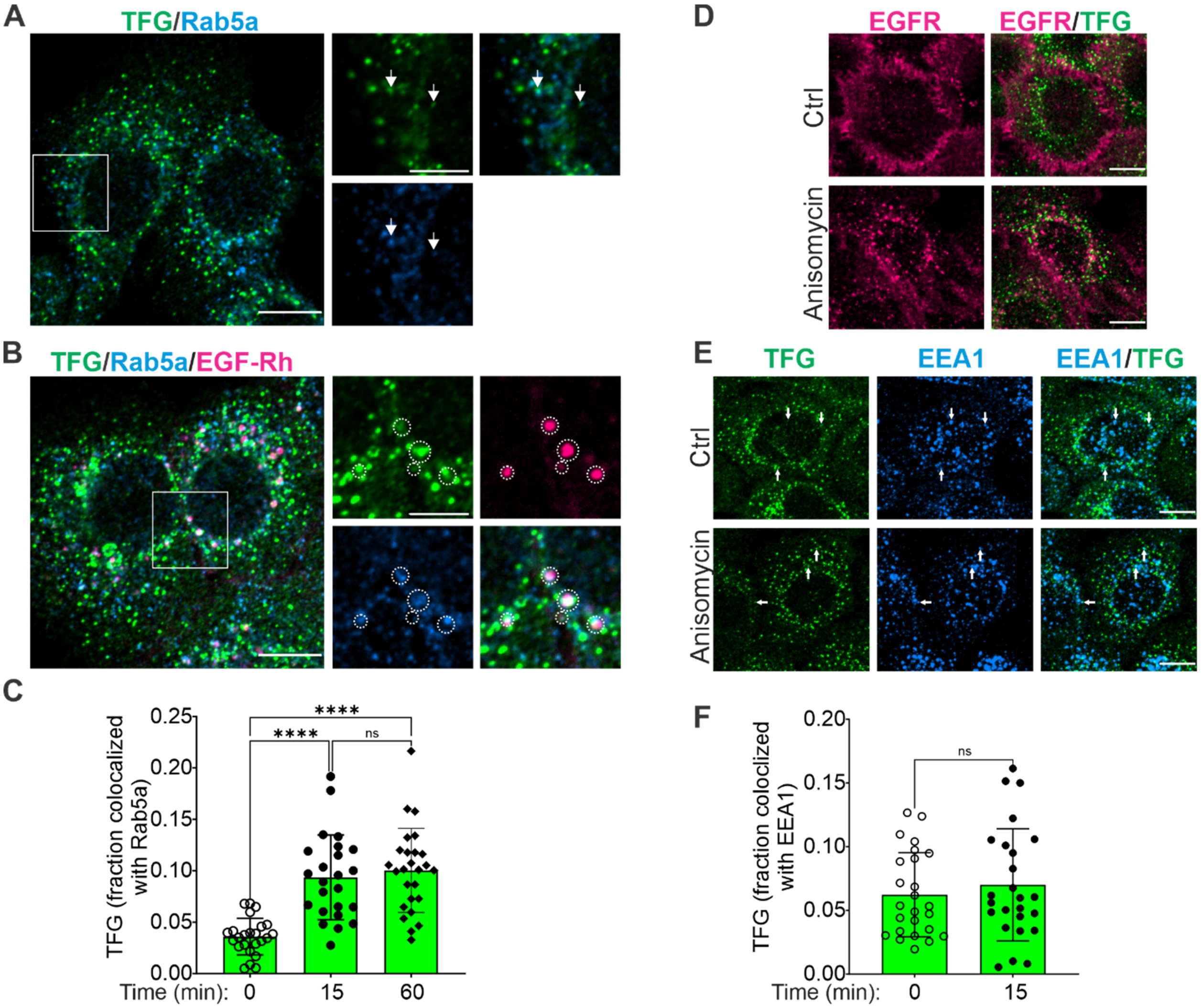


**Figure S2. Localization of TFG with endosomes in EGF- and anisomycin-stimulated cells.**

(**A**-**C**) HCS3 cells were untreated (***A***) or treated with 10 ng/ml EGF-Rh for 15 (***B***) or 60 (images are not shown) min, fixed and stained with TFG and Rab5 antibodies. 3D confocal images were acquired through 640 nm (cyan, Rab5), 561 nm (magenta, EGF-Rh) and 488 nm (green, TFG) channels. Arrows point at examples of TFG localization with Rab5. Circles indicate examples of colocalization of TFG, EGF-Rh and Rab5. Insets represent enlargements of the regions indicated by rectangles. All images are single sections through the middle of the cell of representative 3D images. Scale bars, 10 μm in full images and 5 μm in insets. In **C**, fraction of TFG co-localized with Rab5a is quantified. Bar graph represents mean values with SDs (n of 24-26 FOVs from 3 independent experiments). One-way ANOVA. Tukey's multiple comparison test.

(**D**-**F**) HSC3 cells untreated or treated with 100 nM Anisomycin for 15 min were immunolabeled with TFG and either EGFR (**D**) or EEA1 (**E**) antibodies. 3D confocal images were acquired through 640 nm (magenta EGFR or cyan EEA1) and 488 nm (green, TFG) channels. 3D maximum projections are shown. Scale bars, 10 μm. In **F**, the fraction of TFG co-localized with EEA1 was quantified. Bar graph represents mean values with SDs (n of 28 FOVs from 3 independent experiments). Mann-Whitney test.

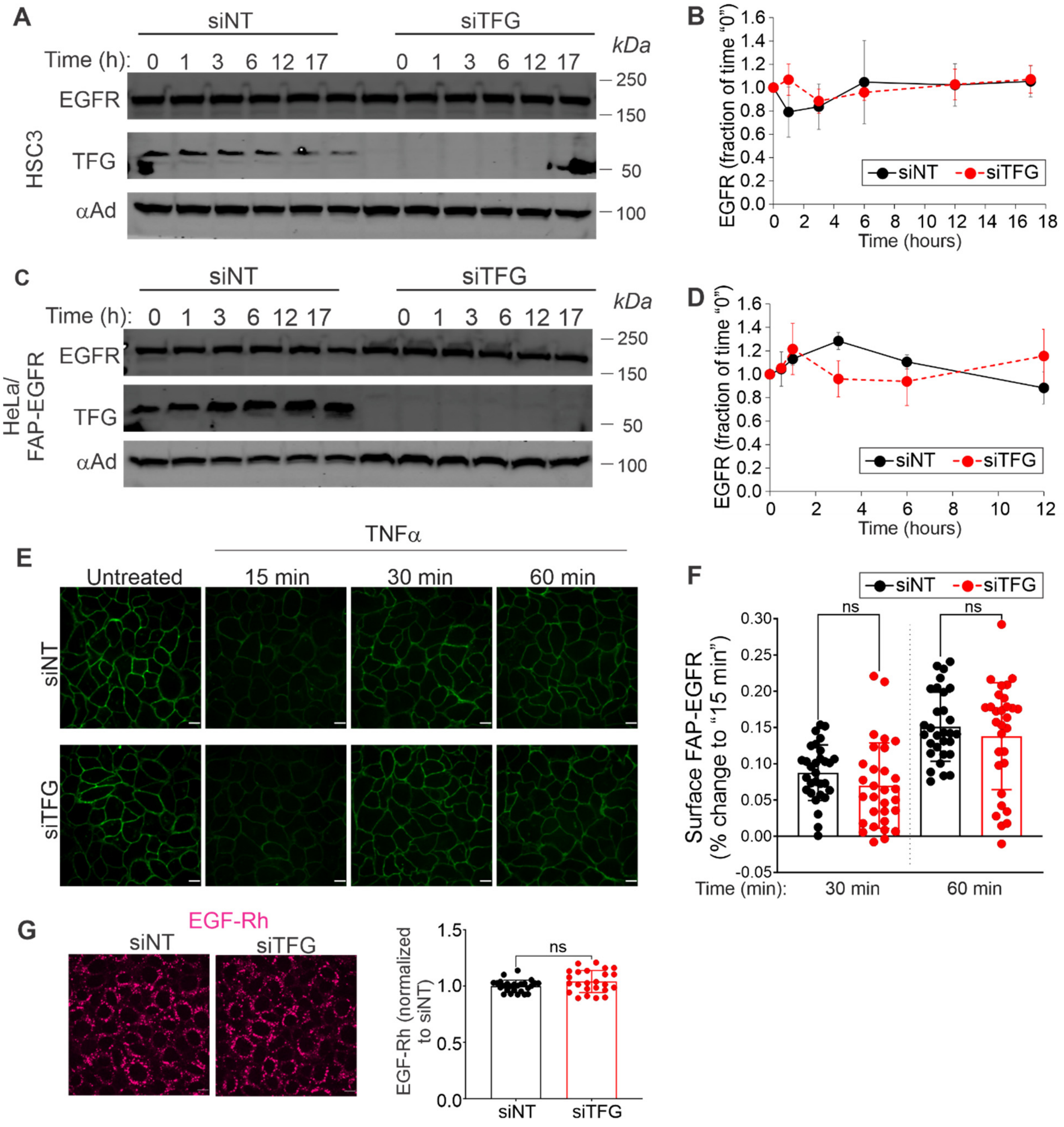


**Figure S3. TFG depletion does not affect constitutive EGFR degradation and recycling of internalized ligand-free EGFR.**

(**A**-**B**) HSC3 or (**C**-**D**) HeLa/FAP-EGFR cells transfected with non-targeting (siNT) or TFG (siTFG) siRNAs were incubated with 20 μg/ml cycloheximide for indicated times. Cells were then lysed, and the lysates were probed by Western blotting with antibodies to EGFR, TFG and α-adaptin (αAd, loading control). The amount of EGFR divided by the amount of α-adaptin was then normalized by this value at time "0" and plotted against time. Graphs show mean values with SEMs of 3 independent experiments.

(**E-F**) HeLa/FAP-EGFR cells transfected with siNT or siTFG were incubated with 10 ng/ml TNFα for indicated times. Internalization was stopped by placing the cells on ice, and surface FAP-EGFR was labeled with MG-B-Tau. Cells were fixed and imaged through the 640 nm channel (*green*, surface EGFR). Representative sections through the middle of the cells are shown. Intensity scales are identical on all images. Scale bars, 10 μm. In ***F***, fluorescence intensities of MG-B-Tau per FOV were quantified. The mean value obtained at 15 min was subtracted from each FOV value at indicated time points and the resulting difference normalized by the mean value from untreated cells to obtain the fraction of fluorescence change (recovery from 15 min). Bar graph shows mean with SDs (n of 30 from 3 independent experiments). Kruskal-Wallis test. Dunn's multiple comparison test.

(**G**) HeLa/FAP-EGFR cells transfected with siNT or siTFG were incubated with 10 ng/ml EGF-Rh for 15 min and fixed. Images were acquired through 561 nm channel (*magenta*, EGF-Rh). Intensity scales are identical on both images. Scale bars, 10 μm. The sum intensity of rhodamine fluorescence per FOV was calculated and normalized by the mean value obtained in siNT. Graph shows mean with SDs (n of 24-26 FOVs from 3 independent experiments). Unpaired t-test.

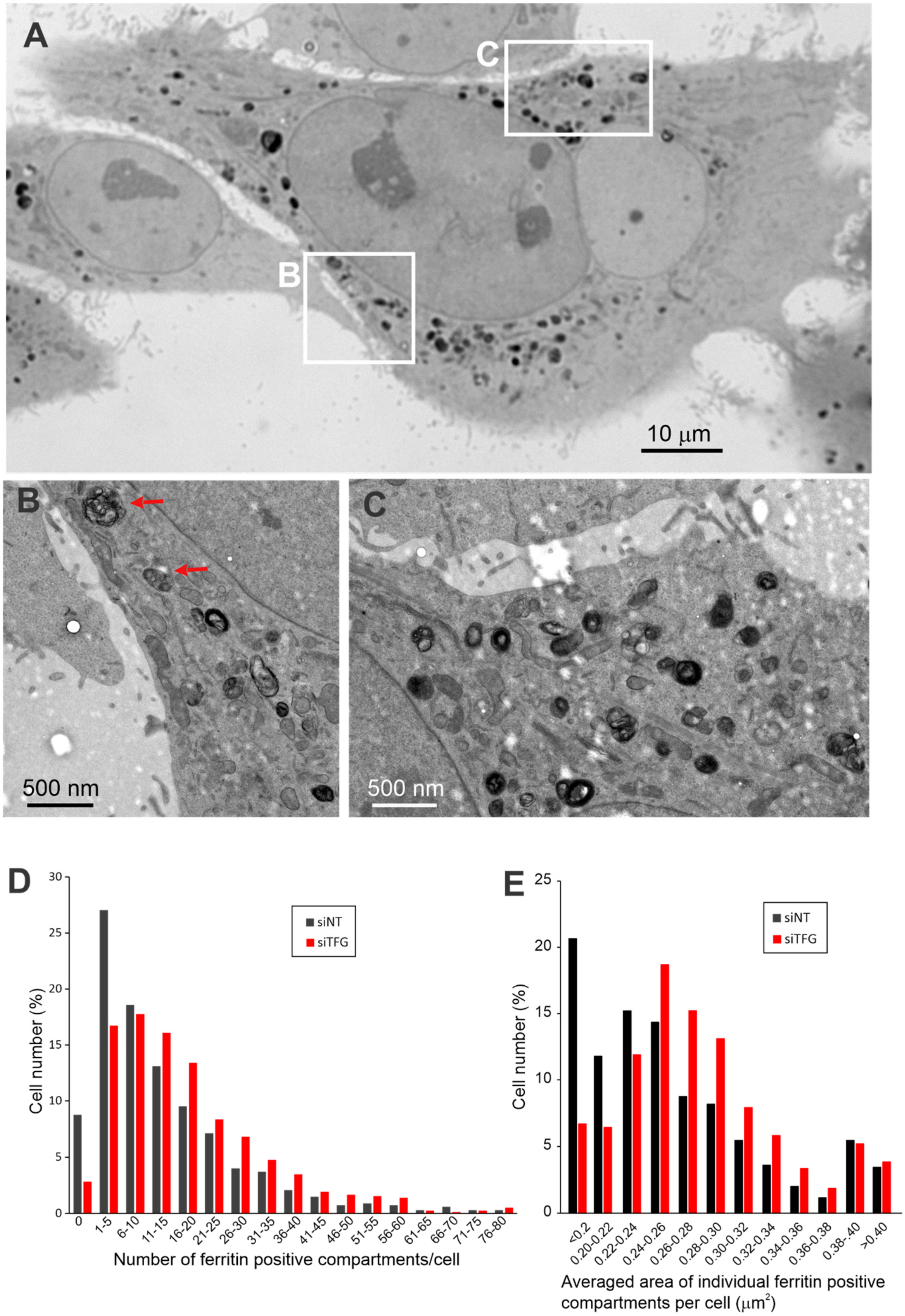

A
C
B
10 μm
B
500 nm
C
500 nm
D
siNT
siTFG
Cell number (%)
Number of ferritin positive compartments/cell
0
1-5
6-10
11-15
16-20
21-25
26-30
31-35
36-40
41-45
46-50
51-55
56-60
61-65
66-70
71-75
76-80
E
siNT
siTFG
Cell number (%)
<0.2
0.20-0.22
0.22-0.24
0.24-0.26
0.26-0.28
0.28-0.30
0.30-0.32
0.32-0.34
0.34-0.36
0.36-0.38
0.38-.40
>0.40
Averaged area of individual ferritin positive compartments per cell (μm²)

**Figure S4. TFG siRNA knockdown increases the number and size of endosomal compartments.**
HSC3 cells were transfected with non-targeting (NT) or TFG siRNA and used for experiments after 3 days. Cells were incubated with 200 nM ferritin for 30 min, fixed and processed for CLEM analysis.

(**A**) Representative light microscopy (LM) image of cells transfected with NT.

(**B** and **C**) EM correlates of semi-serial (within 200 nm) of the regions marked by white rectangles in ***A***. Red arrows show examples of multivesicular bodies.

(**D**) Quantification of the number of ferritin-positive endosomes per cell was performed in in 679 and 808 cells in control and TFG-depleted cells, respectively. The percent distribution of resulting endosome/cell values within cell population is presented.

(**E**) Quantification of the average area of individual ferritin-positive endosomes per cell was performed in 679 and 808 cells in control and TFG-depleted cells, respectively. The percent distribution of resulting values of averaged area of endosomes within cell population is presented.
